# Inhibition of pterygium cell fibrosis by the Rho kinase inhibitor

**DOI:** 10.1101/2024.10.05.616834

**Authors:** Jiannong Dai, Naga Pradeep Rayana, Michael Peng, Chenna Kesavulu Sugali, Devon Hori Harvey, Kamesh Dhamodaran, Eric Yu, Shaohui Liu, Weiming Mao

**Affiliations:** Eugene and Marilyn Glick Eye Institute, Indiana University School of Medicine; Department of Ophthalmology, Indiana University School of Medicine; Fulton Science Academy Private School, Alpharetta, GA; Department of Biochemistry & Molecular Biology, Indiana University School of Medicine; Department of Pharmacology and Toxicology, Indiana University School of Medicine; Stark Neurosciences Research Institute, Indiana University School of Medicine

**Author notes:** **Corresponding author:** Weiming Mao Ph.D., Associate Professor, Jay C. and Lucile L. Kahn Scholar in Glaucoma Research and Education, Showalter Scholar, Eugene and Marilyn Glick Eye Institute, Department of Ophthalmology, Department of Biochemistry & Molecular Biology, Department of Pharmacology and Toxicology, Stark Neurosciences Research Institute, Indiana University School of Medicine, RM305v, 1160 W. Michigan St, Indianapolis, IN, 46202.

**Keywords:** Pterygium, recurrence, Rho kinase inhibitor, fibrosis, health disparity

## Abstract

Pterygium is an ocular disease in which the conjunctival tissue invades the cornea. When the pterygium tissue reaches the pupillary region, the visual function of the patient is affected. Currently, surgical removal is the only effective treatment. However, the recurrence rate of pterygium after surgery can be high. Pterygium is also a health disparity issue since it is more prevalent in the Hispanic and Latino American population. In this study, we determined if the Rho kinase inhibitor can be used to prevent pterygium recurrence since its anti-fibrosis effects have been reported in other cell and tissue types. We cultured primary pterygium cells from pterygium tissues from Hispanic and Latino American, African American, Caucasian, and Asian donors, and used those cells for viability assays, scratch assays, migration assays, and immunostaining of F-actin, fibronectin, collagen I and α smooth muscle actin. We found that the Rho kinase inhibitor Y27632 decreased cell viability, wound healing, cell migration, as well as the expression of extracellular matrix and myofibroblast markers in cultured pterygium cells. We believe that Rho kinases inhibitors are a potential post-surgical treatment to prevent pterygium recurrence.

## Introduction

Pterygium is an ocular disease in which the conjunctival tissue grows onto the cornea. When the pterygium tissue reaches the pupillary region and blocks the visual axis, the visual function of the patient is affected. The pterygium tissue appears to be triangular/wing-shaped with the apex/head toward the center of the cornea and the body close to the limbus. Currently, surgical removal is the only effective treatment of pterygium. However, the recurrence rate of pterygium after surgery can be as high as 89% ^1,2^. Therefore, it is important to develop an effective medicine to prevent pterygium recurrence.

The etiology of pterygium outgrowth is not clearly understood. Pterygium tissues mainly consist of epithelial and fibrovascular tissues. During the outgrowth of pterygium, inflammatory cell infiltration and extracellular matrix accumulation play important roles ^3,4^. In normal eyes, the limbal region contains stem cells, and this region functions as a barrier which prevents abnormal epithelial tissue ingrowth ^5,6^. In contrast, if the limbal stems cells and/or their environment are compromised, the neighboring conjunctival tissue will encroach onto the cornea ^5,6^. The primary risk factor of pterygium is UV exposure, which may induce oxidative stress and DNA damage which further disrupt the balance among proinflammatory cytokines, apoptotic factors, and growth factors ^1,5,7,8^. Accordingly, pterygium are more prevalent in the equatorial region ^9^. Ethnic group wise, Hispanic and Latino Americans have a high risk in developing pterygium ^1,10^.

Although pterygium removal is safe and effective, after pterygium removal, excessive fibrosis frequently leads to pterygium recurrence. The current method to decrease pterygium recurrence includes surgical techniques such as the bare sclera technique (recurrence up to ∼89%), conjunctival autograft (∼3-33%), amniotic membrane transplantation (∼4-41%), conjunctival transpositional flap (∼3-33%), as well as antimitotic medicines such as mitomycin C (∼4-9%) and 5-fluorouracil (∼11-60%) ^1^.

Since pterygium recurrence involves fibrosis, we believe that a novel group of medicine, the Rho kinase inhibitor, may have the potential in preventing pterygium recurrence. The Rho kinase inhibitor inhibits the Rho-ROCK signaling pathway which plays an important role in fibrotic events including cell adhesion, actomyosin cytoskeleton organization, cell adhesion, and extracellular matrix production ^11^. Igarashi et.al reported that the upregulation of sphingosine-1 phosphate (S1P), an activator of the Rho-ROCK signaling pathway, promotes fibrosis in human pterygium cell cultures ^12^. Recently, Xie and colleagues showed that the Rho kinase inhibitor Y27632 inhibits TGFβ1-induced fibrosis in human pterygium fibroblast cells ^13^. However, whether Y27632 inhibits fibrosis in human pterygium cells under “natural”/unstimulated conditions is unclear. Also, since the study conducted by Xie and colleagues used samples from the Chinese population, it is not clear whether such inhibition exists in non-East Asian populations while health disparity is an emerging issue. Therefore, we conducted this study to answer these questions.

## Materials and methods

### Pterygium tissue collection

Pterygium tissues were collected from pterygium surgeries performed by a board-certified ophthalmologist at Eugene and Marylin Glick Eye Institute at Indiana University. The collection of pterygium tissues was conducted according to Declaration of Helsinki principles. Human subjects were provided written informed consent. This study was approved by the Institutional Review Board of Indiana University–Purdue University at Indianapolis (approval number 2009784394). The donors included Hispanic and Latino Americans (6), African Americans (3), Caucasians (2), and Asian (1).

### F-actin staining

Primary human pterygium cells were grown on a 96-well cell culture plate and treated with either DMSO as a control or 100 µM Y27632 for 3 or 5 hours. At the end of treatment, the cells were fixed with 4% paraformaldehyde in PBS for 30 minutes at 4°C and treated with 0.5% Triton X-100 for 10 minutes. The cells were then incubated with Phalloidin-Alexa 488 (Thermo Fisher Scientific, Waltham, MA) at 3.3 pM overnight (∼16 hours) at 4°C and counter stained with DAPI. The cells were imaged using the Nikon Ti2 inverted microscope and the Qi2 digital camera (Nikon Instruments Inc. Melville, NY).

### Pterygium cell proliferation and culture

Pterygium tissues were used to establish primary human pterygium cell strains. Some tissues were dissected into small sections about 2-5mm in diameter/length. Individual tissue section was placed in the 60mm cell culture dish containing the Opti-Modified Eagle’s Minimum Essential Medium (Opti-MEM) (Thermo Fisher Scientific, Waltham, MA) supplemented with 5% or 10% fetal bovine serum, 1% penicillin and streptomycin, and 2 mmol/L glutamine. The tissue was incubated in the cell culture incubator at 37°C with 5% CO2. Medium was changed every other day. Cell outgrowth from the tissue was monitored. When cells were close to confluent, they were passaged using TrypLE (Thermo Fisher Scientific). The cells within 4-6 passages were used for all the experiments. Due to our culture methods, the pterygium cells that we used contained both epithelial cells and fibroblast cells. ^14^

### Cell viability assays

Primary human pterygium cells were seeded on a 96-well cell culture plate (2000 cells/well). The next day, the cells were treated with either DMSO as a control, or mitomycin C or Y27632 at different concentrations for 48 hours, and then used for MTT (Sigma-Aldrich Saint Louis, MO) 5mg/ml in PBS or XTT assays (Thermo Fisher Scientific) following manufacturer’s instructions. Absorbance was measured using Synergy H1 Hybrid Multi-Mode plate reader (BioTek Instruments, Winooski, VT).

### Compounds

The Rho kinase inhibitor, Y27632 was purchased from Tocris Bioscience (Minneapolis, MN; Catalog #:1254). Mitomycin C was purchased from Sigma (Catalog #: M4287). Both compounds were dissolved in dimethyl sulfoxide (DMSO) to prepare a stock solution at the concentration of 10 mM.

### Scratch assays

Primary pterygium cells were seeded in 12-well to 24-well culture plates at a density of 2 × 10^5^/cm^2^. After the cells became confluent, a scratch was made using a 1ml pipette tip in each well, and the detached cells were removed using PBS wash. After replacing PBS with culture medium, the cells were treated with either 0.1% DMSO or 10µM Y27632.

Some cells cultured in 12-well plates were scratched using a 5.7 mm wide plastic spatula. The cells were treated with DMSO or Y27632. Some cells were initially treated with Y27632 for 3 days followed by Y27632 removal.

Images showing the area of the scratch were taken at different time points using the Nikon Ti2 inverted microscope with the Nikon Advanced Modulation Contrast. The images were analyzed using the ImageJ software (National Institute of Health, Bethesda, MD). After the scale was set (“Analyze”➔ “Set scale”), the area of scratch was outlined using the “Free hand selection” tool and then measured using the “Measure” function (“Analyze”➔ “Measure”). The area of the initial scratch (A_0_) was set at 100%, and the area in the following days (A) was compared to that of the initial scratch (A/A_0_%).

### Cell migration from pterygium explants

The pterygium tissues obtained from the patient were cut into 3 equal pieces. Each piece was further cut into 2 halves and seeded in the 12 well cell culture plate (Thermo Fisher Scientific) and incubated with Opti-MEM supplemented with FBS and antibiotics. One half was treated with DMSO as a control while the other half was treated with 10µM of Y27632. Medium and treatment were carefully changed every other day. Images of the tissues were taken using the Nikon Ti2 inverted microscope (Nikon Instrument, Melville, NY) daily. The area of the cells that migrated and proliferated from the tissue explant was measured using the ImageJ software.

### Immunostaining

Human pterygium cells were seeded in a 12 well glass bottomed plate (1×10^5 cells/well). Cells were incubated overnight before they were treated with DMSO as a control or 10 µM Y27632 for 4 days. After treatment, the cells were fixed with 100% methanol (Thermo Fisher Scientific) for 20 minutes at −20°C. Fixed cells were washed with PBS and blocked using the Superblock TW blocking buffer (Thermo Fisher Scientific) in PBS for 60 minutes at room temperature. The cells were then incubated with the primary antibody, mouse anti-L-Smooth muscle actin (1:200; catalog#: F3777; Sigma-Aldrich, St. Louis, MO) or rabbit anti-fibronectin antibody (1:200; catalog#: ab1945; Sigma-Aldrich) or rabbit anti-collagen-1 antibody (1:200; catalog#: ab215969; Abcam, Boston, MA) overnight at 4°C. After PBS wash, the cells were incubated with the secondary antibody (goat anti-mouse/-rabbit Alexa 488 or 568; 1:200; Thermo Fisher Scientific) for 2 hours at room temperature. The nuclei were counter stained with DAPI (Thermo Fisher Scientific). The cells were visualized under Nikon Eclipse Ti2 inverted microscope (Nikon).

### Statistical Analysis

The data were analyzed using Prism Graphpad 9.5.1 (GraphPad Prism Software Inc., San Diego, CA) for one-way ANOVA and posthoc tests or Student’s t-tests. P values less than 0.05 were considered statistically significant.

## Results

### The Rho kinase inhibitor Y27632 decreased pterygium cell viability

Ideally, to prevent pterygium recurrence, the compound should be able to decrease cell viability but not cause excessive cell damage which may be toxic to other ocular surface tissues. Therefore, we determined the effect of Y27632 on pterygium cell viability.

We first determined the dose-response effect of Y27632 and compared it to that of mitomycin C, which is often used clinically during pterygium surgeries to decrease recurrence ^2^. We seeded pterygium cells onto 96-well plates and treated them with 0.1% DMSO (as a control), different concentrations of mitomycin C or Y27632 for 2 days (Figure 1) before viability assays. We found that 10µM Y27632 (a frequently used concentration in eye research), had similar effect on reducing pterygium cell viability as 10µM mitomycin C.

**Figure 1.**
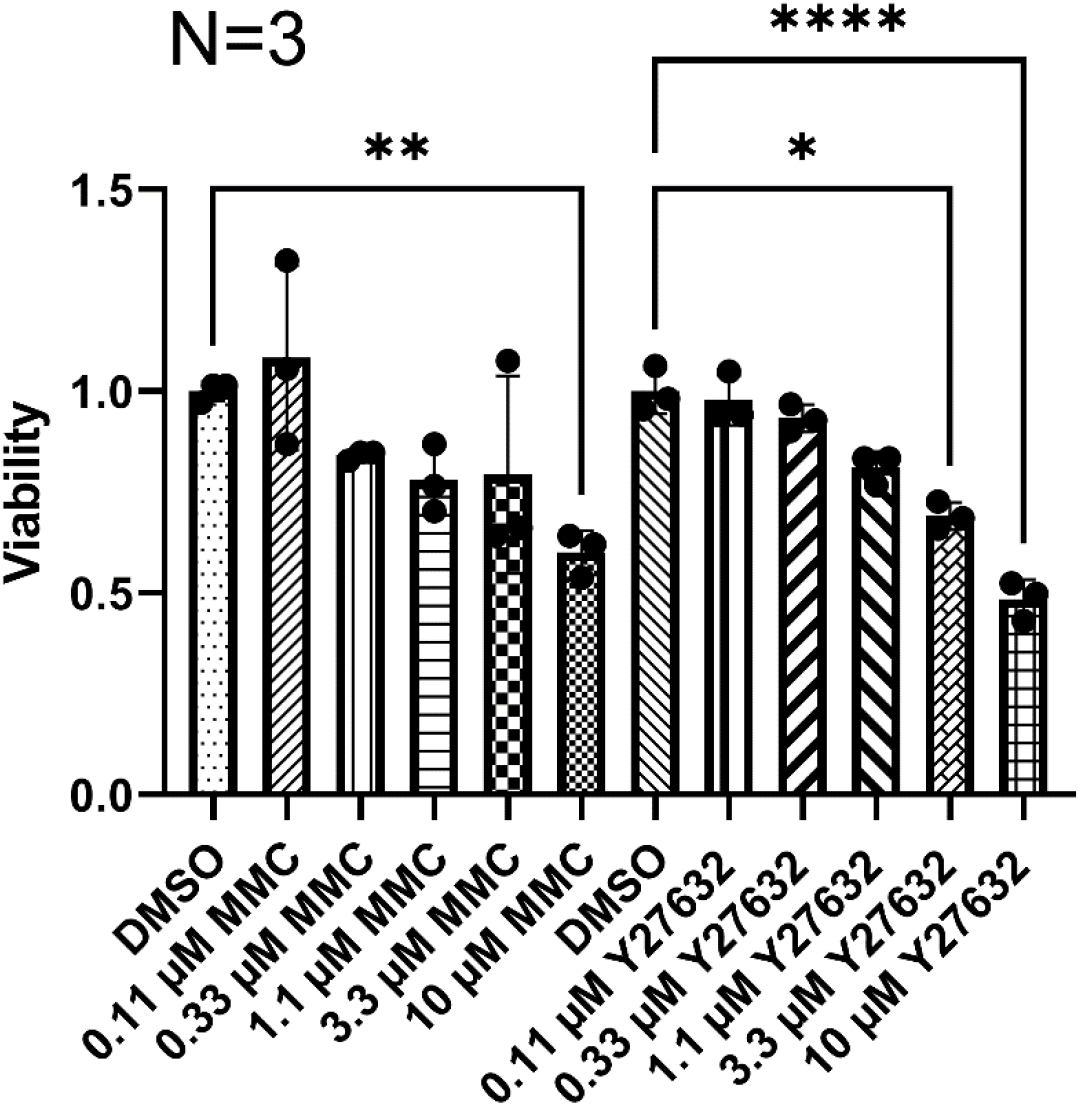
The Rho kinase inhibitor Y27632 dose-dependently decreased pterygium cell viability. Primary pterygium cells were treated with 0.1% DMSO or mitomycin C (MMC) or Y27632 at indicated concentrations for 2 days before they were used for MTT or XTT assays. Columns and error bars: means and standard deviations. One-way ANOVA and posthoc tests were used for statistical analysis. N=3. *: p<0.05, **: p<0.01, ****: p<0.0001.

We then focused on 10µM Y27632 and repeated viability assays in 4 additional pterygium cell strains (Figure 2). Similarly, Y27632 significantly decreased viability (p< 0.05) in all 4 cell strains. However, we also observed that the response to Y27632 varied among these cell strains (56%, 14%, 16%, and 21% decrease in viability in Figure 2A-D, respectively).

**Figure 2.**
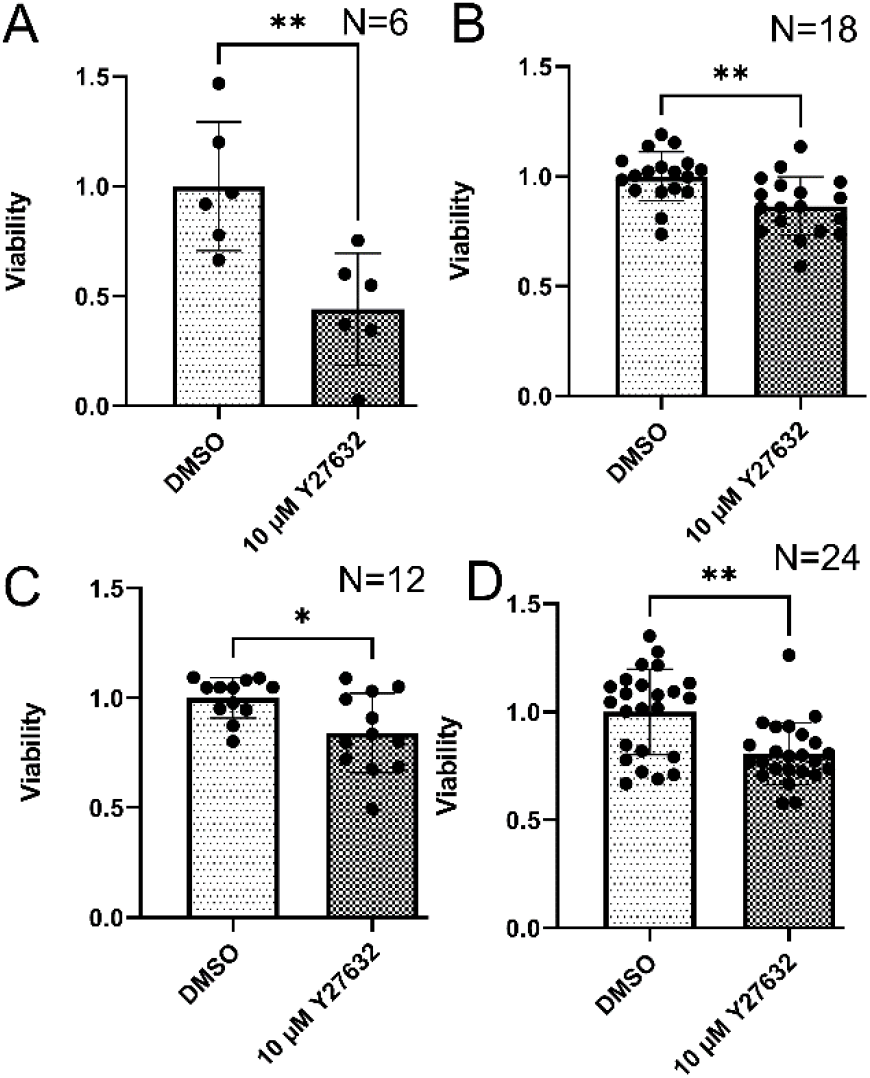
The Rho kinase inhibitor Y27632 decreased pterygium cell viability in multiple cell strains. Four different primary pterygium cell strains (A-D) were treated with 0.1% DMSO or 10µM Y27632 for 2 days before they were used for MTT or XTT assays. Columns and error bars: means and standard deviations. Student’s t-test was used for statistical analysis. Sample sizes (N) are indicated in each panel. *: p<0.05, **: p<0.01.

### The Rho kinase inhibitor Y27632 inhibited wound healing

To further determine the effect of Y27632 on wound healing, we conducted scratch assays using primary pterygium cells.

Pterygium cells with passage numbers less than 4 were seeded onto multi-well culture plates. When the cells were confluent, scratches were made using 1ml tips. The acellular area created by scratch was imaged and measured. The scratch assay showed a decrease in cell coverage by 10µM Y27632 compared to 0.1% DMSO, and 4 representative strains are shown in Figure 3A-D.

**Figure 3.**
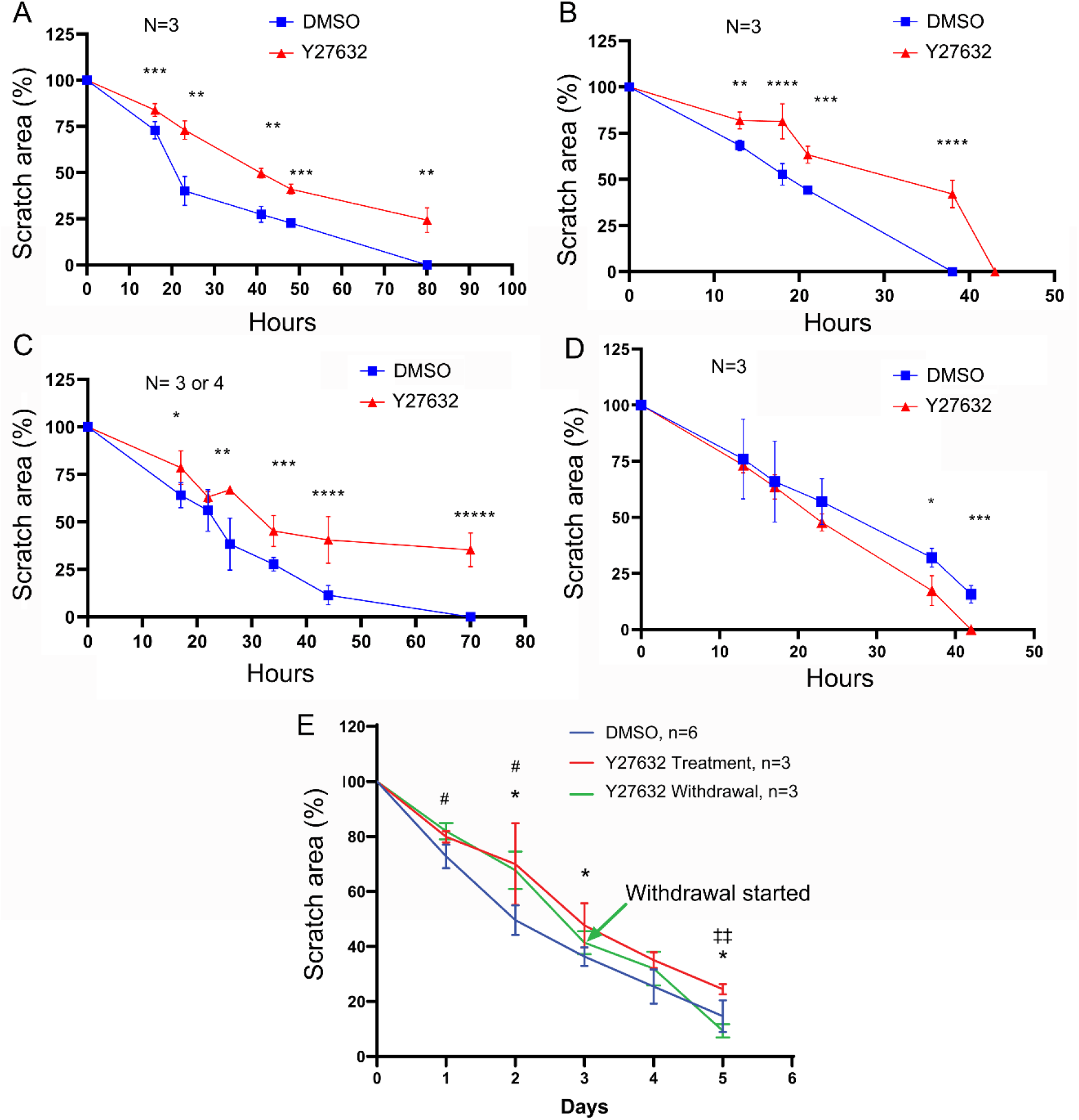
The Rho kinase inhibitor Y27632 inhibited wound healing. Four different primary pterygium cell strains (A-D) were used for scratch assays. After uniform scratches were created in confluent cell cultures, the cells were treated with 0.1% DMSO (Blue) or 10µM Y27632 (Red). In another cell strain (E), wide scratches (5.7mm) were created, and the cells were treated with DMSO (blue), Y27632 (red), or Y27632 for 3 days followed by Y27632 withdrawal. Scratch areas (the area of the acellular region) were measured, and means and standard deviations (error bars) were plotted over time. Student’s t-test was used for statistical analysis for individual time point. (A-D): *: p<0.05, **: p<0.01, ***: p<0.001, ****: p<0.0001, *****: p<0.00001. (E): *:DMSO vs. Y27632 withdrawal, p<0.05; *: DMSO vs. Y27632, p<0.05; ‡‡: Y27632 vs. Y27632 withdrawal, p<0.01.

To determine whether the inhibition of Y27632 is reversible, we cultured another cell strain and created wide scratches (5.7mm) to allow longer observation windows before the scratches are fully covered by the cells. The cells were treated with 0.1% DMSO, 10µM Y27632, or 10µM Y27632 for 3 days followed by Y27632 withdrawal (Figure 3E and supplementary Figure S1). We found that Y27632 significantly delayed wound healing (days 1-3; p<0.05). However, once Y27632 was removed (Y27632 withdrawal), the scratches in these cells were smaller than Y27632 treated cells (Y27632 treatment) but larger than DMSO treated cells (DMSO) on day 5. Our data suggest that 1) the inhibition of wound healing by Y27632 is reversible; and 2) the “residual” effect of Y27632 can last for at least 2 days.

### The Rho kinase inhibitor Y27632 inhibited cell migration from pterygium explants

We determined if Y27632 was able to inhibit cell migration from pterygium tissues since this process is the key to pterygium recurrence.

We placed pterygium explants (collected from pterygium surgeries) on culture plates with culture medium, and treatment them with 0.1% DMSO as a control or 10µM of Y27632 (sample from two different donors are shown in Figure 4A and B). Cell migration and/or proliferation was monitored by measuring the area that the outgrowing pterygium cells covered. We found that Y27632 slowed down cell migration and/or proliferation from both groups of pterygium explants. We also studied cell migration in 10 additional explants. However, those explants detached within one or two days after Y27632 treatment (DMSO treated explant did not detach), and we were not able to obtain quantitative migration assay data from those explants. Nevertheless, their early detachment indicates an almost complete inhibition of cell migration.

**Figure 4.**
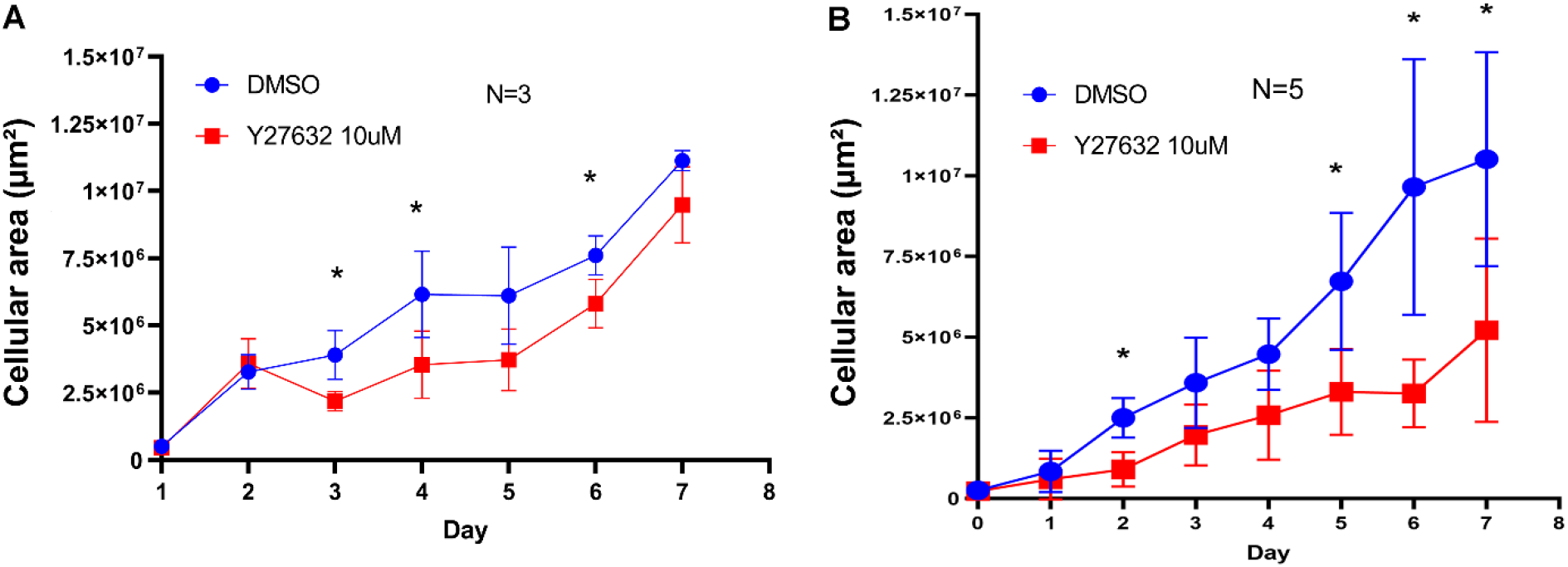
The Rho kinase inhibitor Y27632 decreased cell migration. Pterygium tissues from two different donors (A and B) were collected from pterygium surgeries. For the tissues from each patient, 3 pieces of tissues were dissected. Each piece of tissue was further cut into two halves, and both halves were placed onto culture dishes. One half tissue was treated with 0.1% DMSO (Blue), while the other half was treated with 10µM Y27632 (Red). Cellular areas (the area that the migrating pterygium cells covered) were measured, and means and standard deviations (error bars) were plotted over time. Student’s t-test was used for statistical analysis for individual time point. N=3 or 5. *: p<0.05.

### The Rho kinase inhibitor inhibited F-actin and fibrosis-related proteins in pterygium cells

We used immunofluorescence to determine the effect of Y27632 on F-actin (Figure 5), extracellular matrix, and α-smooth muscle actin (α-SMA) of pterygium cells (Figure 6). We found that actin stress fibers (F-actin), as well as fibronectin and collagen type 1, two of the key proteins for wound healing, were decreased by treatment with 10µM Y27632. L-SMA, the myofibroblast marker, was also inhibited by Y27632.

**Figure 5.**
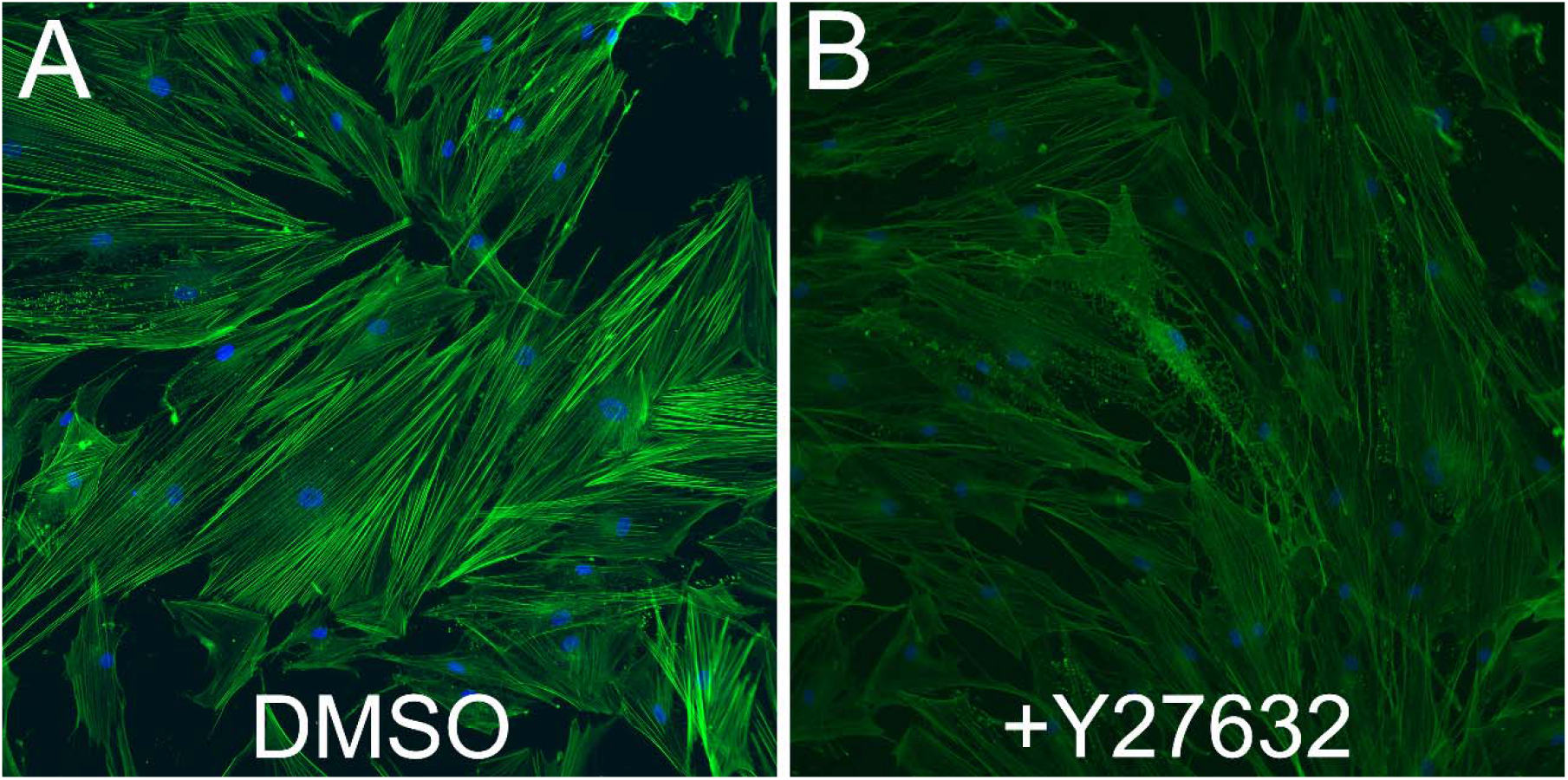
The Rho kinase inhibitor Y27632 inhibited actin polymerization in primary pterygium cells. Primary pterygium cells were treated with 0.1% DMSO (A) or 10µM Y27632, fixed and stained with phalloidin-Alexa 488 for F-actin (green) and counterstained with DAPI (blue) for nuclei.

**Figure 6.**
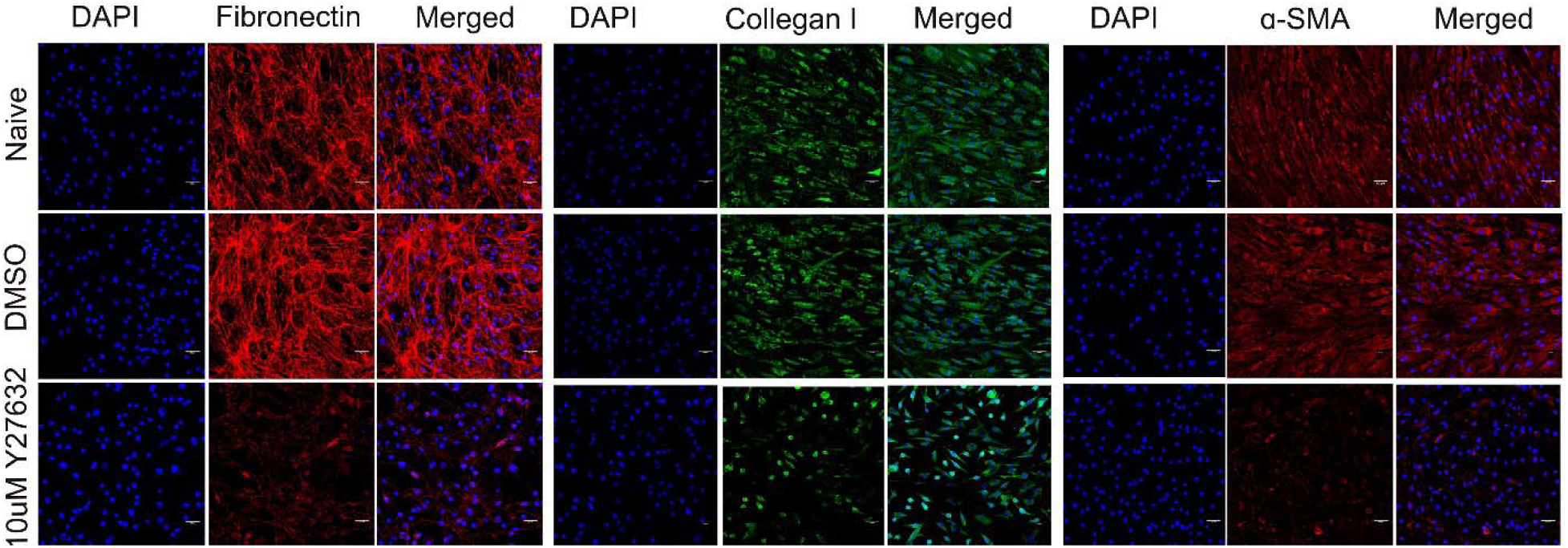
The Rho kinase inhibitor Y27632 inhibited fibronectin, collagen I and α-smooth muscle actin (α-SMA) in primary pterygium cells. Primary pterygium cells were treated with indicated agents and used for immunostaining. DAPI was used for nuclear staining. Three different strains were used and representative images of one strain are shown.

## Discussion

The major challenge of pterygium surgery is recurrence. Our data suggested that the Rho kinase inhibitor Y27632 can decrease cell proliferation and migration, epithelial mesenchymal transition, and extracellular matrix accumulation in unstimulated pterygium cells from non-East Asian donors.

As described previously, Rho kinase inhibitors inhibit fibrosis in multiple ocular tissues. However, they may also promote skin wound closure^15^, corneal endothelial cell adhesion and proliferation ^16^. We believe that these seemingly contradictory effects are likely to be cell/tissue type dependent.

Currently, there have been no post-surgical treatment available for the prevention of pterygium recurrence (mitomycin C and 5-fluorouracil are used during surgery). Our studies showed that the Rho kinase inhibitor Y27632 is a promising treatment. We believe that Rho kinase inhibitors, if can be used for pterygium recurrence prevention clinically, have two advantages:

1. Feasibility. Some Rho kinase inhibitors are already FDA-approved for ocular use for treating glaucoma ^17^. Therefore, ophthalmologists may have the opportunity to use it “off-label” once more supporting data are available.
2. Safety. FDA approved Rho kinase inhibitors have not shown significant toxicity to ocular surface tissues ^18,19^. Instead, they only induce minor side effects such as hyperemia, blepharitis, ocular irritation, and increased lacrimation which are usually tolerable ^19^. Compared to mitomycin C and 5-fluorouracil, which have high cellular toxicity and are both mutagens, Rho kinase inhibitors are considered safe.
3. Convenience. Topical Rho kinase inhibitors can be self-administrated by the patient.

However, there are still some key questions and potential issues that need to be addressed before Rho kinase inhibitors can be used clinically:

1. The effect of Rho kinase inhibitors in vivo.

Our current data are all based on cell cultures. The results that we observed in vitro do not necessarily guarantee the same effect in vivo. Therefore, in vivo studies, such as using rodent pterygium research models, are needed with the clinically approved the Rho kinase inhibitors.

2. Hyperemia

Hyperemia is a common side effect of Rho kinase inhibitors due to vessel dilation ^19^. It is generally believed that hyperemia may promote fibrosis by providing nutrition, oxygen, and proinflammatory factors to the wound area. Therefore, the anti-fibrotic effect of Rho kinase inhibitors may be counteracted or even surpassed by hyperemia. Our in vitro system lacks blood vessels, and therefore cannot be used to address this possibility. Again, this question can only be answered by using in vivo models. The other side effects of Rho kinase inhibitors, such as blepharitis, are unlikely to promote fibrosis and therefore not discussed here.

In summary, we found that Rho kinase inhibitors may be a potential post-surgical treatment to minimize pterygium recurrence.

## Acknowledgments

This study was supported by the National Institute of Health/National Eye Institute Award Numbers R01EY026962 (WM), R01EY031700 (WM), R21EY033929 (WM), BrightFocus Foundation G2023009S (WM), and a challenge grant from Research to Prevent Blindness (Department of Ophthalmology, Indiana University School of Medicine). The content is solely the responsibility of the authors and does not necessarily represent the official views of the National Institutes of Health.

## Author contributions

SL and WM conceived the idea and designed the experiments; SL provided clinical samples. JD, NPR, MP, and DHH conducted the experiments. JD, NPR, MP, CKS, KD, EY, and WM analyzed the data. JD, NPR, MP, CKS, DHH, KD, and WM wrote the manuscript.

## Data availability

The raw data are available upon request. Please send such request to WM.

