## Supplemental Figure 1 legend for "Inhibition of pterygium cell fibrosis by the Rho kinase inhibitor"

Supplemental Figure 1. Representative images of scratch assay.

Pterygium cells were treated with DMSO (A-C), Y27632 (D-F), or Y27632 for only 3 days (G-I). The black lines were makers made at the bottom of the culture plate to indicate the approximate location of the scratch to facilitate serial imaging over time.
