## Supplementary figures and images for "Inhibition of pterygium cell fibrosis by the Rho kinase inhibitor"

### Supplemental Figure 1

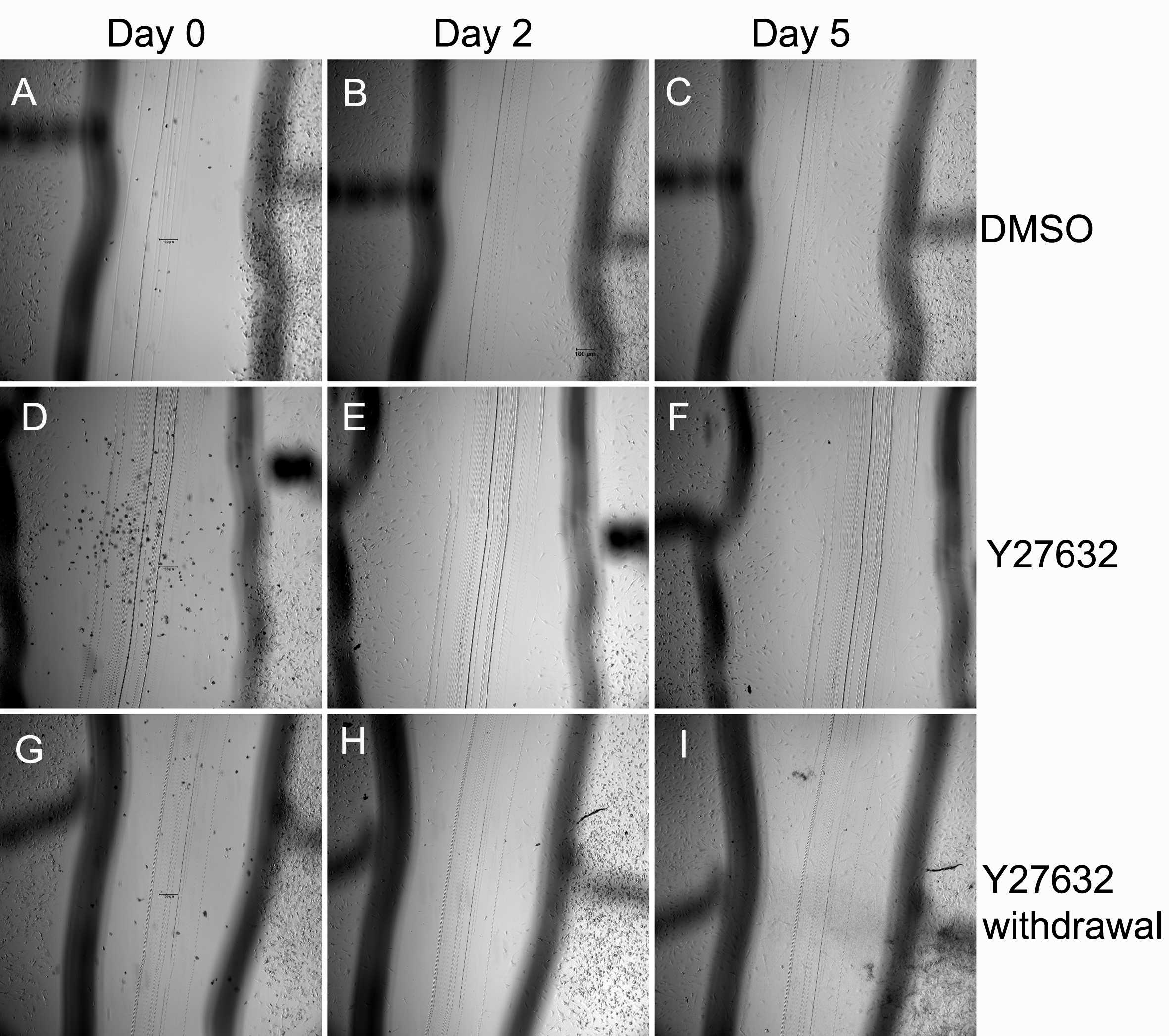
